# Development of Novel NPC1-Targeting Autophagy Inhibitors for the Treatment of Uveal Melanoma

**DOI:** 10.64898/2026.11.12.766269

**Authors:** Yudai Kudo, Sarah Scoles, Jia Yu Ning, Geyi Li, Michael J. Heiferman, Satoshi Endo, Michael T. Flavin, Kaori H. Yamada

**Affiliations:** Department of Pharmacology & Regenerative Medicine, University of Illinois College of Medicine, Chicago, IL 60612, USA; United Graduate School of Drug Discovery and Medical Information Sciences, Gifu University, 1-1 Yanagido, Gifu 501-1194, Japan; Department of Ophthalmology & Visual Sciences, University of Illinois College of Medicine, Chicago, IL 60612, USA; Center for One Medicine Innovative Translational Research (COMIT), Gifu University, Gifu, 501–1193, Japan; Preemptive Food Research Center, Gifu University, Gifu, 501-1193, Japan; University of Illinois Cancer Center, Chicago, IL 60612, USA

**Keywords:** Autophagy, betulinic acid, cholesterol, lysosome, NPC1, uveal melanoma

## Abstract

**Background:** Uveal melanoma (UM) is the most common primary intraocular malignancy in adults, and up to 50% of patients develop liver metastases. Metastatic UM responds poorly to conventional chemotherapy and currently available immunotherapies, highlighting the need for new therapeutic strategies. Because UM cells rely on lysosome-dependent stress-response pathways, including autophagy, we evaluated previously reported betulinic acid (BetA) derivatives and identified ALS-997 as a potent inhibitor of autophagic flux in UM cells.

**Methods:** The anticancer activity of BetA derivatives was evaluated using the NCI-60 cancer cell line panel and WST-1 viability assays in UM cell lines and normal human melanocytes. Effects on autophagy and lysosomal function were assessed by western blotting for mTOR and autophagy associated proteins, LysoTracker Green imaging, and filipin III/LAMP2 staining. Apoptosis was evaluated by PARP cleavage and caspase-3 cleavage. The interaction of ALS-997 with Niemann–Pick disease type C1 (NPC1) was examined using a cellular thermal shift assay and computational molecular docking. NPC1 expression in UM cells and normal melanocytes was compared by quatitative PCR and western blotting.

**Results:** ALS-997 preferentially reduced the viability of UM cell relative to normal melanocytes. Treatment with ALS-997 disrupted autophagic flux and lysosomal function, induced lysosomal cholesterol accumulation, and activated apoptotic signaling. ALS-997 also altered the thermal stability of NPC1, supporting cellular target engagement, while molecular docking predicted a direct interaction between ALS-997 and NPC1. NPC1 expression was higher in UM cells than in normal melanocytes.

**Conclusions:** ALS-997 is a BetA-derived compound that disrupts lysosomal cholesterol homeostasis and autophagic flux, leading to apoptosis in UM cells. These findings support NPC1 as a potential therapeutic target in UM.

## Background

Uveal melanoma (UM) is the most common primary intraocular malignancy in adults, arising from melanocytes of the uveal tract, which includes the iris, ciliary body, and choroid. It accounts for approximately 3-5% of all melanoma cases and has an incidence of about 5 cases per million annually in the United States. Primary tumors most commonly occur in the choroid, accounting for approximately 90%, and less frequently in the ciliary body or iris ^1,2^. Despite effective local control of the primary tumor with radiotherapy or enucleation, up to 50% of patients develop distant metastases. Most frequently metastasis is to the liver which is observed in approximately 80-90% of metastatic UM cases, even many years after initial diagnosis ^3,4^. Patients diagnosed with metastatic UM have an extremely poor prognosis, with reported median survival periods as short as 3.9 months ^5^. Although tebentafusp, the first FDA-approved therapy for metastatic UM, improves overall survival ^6–11^, its use is restricted to patients with HLA-A*02:01 ^12^. Another FDA-approved treatment for metastatic UM is a liver-directed melphalan ^13,14^, which is limited to patients with liver-predominant disease and is an invasive procedure requiring a specialized center for administration. Despite these advances, eligibility restrictions, treatment burden, toxicity, and limited applicability across all patients leave a critical unmet need to develop treatment of metastatic UM.

At the molecular level, UM is characterized by highly recurrent activating mutations in the G protein subunit alpha q (GNAQ) and G protein subunit alpha 11 (GNA11) genes, which encode paralogous G-protein α subunits, and are found in approximately 80-90% of cases. These mutations activate the mitogen-activated protein kinase (MAPK) pathway and other downstream oncogenic signaling networks. They also promote the activity of the transcriptional regulators Yes-associated protein 1 (YAP) and transcriptional coactivator with PDZ-binding motif (TAZ) ^15^. Additional genetic alterations, such as loss of the BRCA1-associated protein 1 (BAP1) tumor suppressor, are strongly associated with metastatic progression and poor clinical outcomes, making them important prognostic markers in UM ^16–18^. Despite extensive efforts to target these oncogenic pathways therapeutically, direct inhibition has shown limited success in clinical settings, partly due to the complex signaling networks and the relatively low overall mutational burden compared to other melanoma subtypes.

It has been reported that UM cells rely heavily on adaptive stress-response mechanisms, particularly autophagy, to survive under metabolic, oxidative, and therapeutic stress conditions ^19,20^. Autophagy contributes to cancer development and metastasis, but its effects vary depending on tumor type and disease stage ^21^. Recent transcriptomic analyses suggest that autophagy-related gene expression correlates with tumor progression and poor prognosis in UM. Moreover, increased autophagic activity has been observed in UM cell models under stress, supporting a role for autophagy in promoting tumor cell survival and therapeutic resistance ^22–24^.

Because autophagy is a dynamic process rather than a static endpoint, changes in autophagy markers must be interpreted in the context of both autophagosome formation and lysosome-dependent degradation. Elevated lipidated form of microtubule-associated protein 1 light chain 3 (LC3-II) may reflect either enhanced autophagosome formation or impaired autophagosome degradation. Concomitant accumulation of sequestosome 1 (SQSTM1; also known as p62) is often consistent with defective autophagic flux rather than productive autophagy induction ^25^. Thus, therapeutic targeting of the autophagy pathway may involve not only suppressing autophagy initiation but also disrupting lysosome-dependent autophagic flux.

Betulinic acid (BetA) is a naturally occurring pentacyclic triterpene isolated primarily from the bark of birch trees and has attracted considerable attention due to its broad therapeutic and anticancer potential ^26,27^. Extensive preclinical studies have demonstrated that BetA exerts antitumor activity across a wide range of solid malignancies by suppressing tumor growth, inducing apoptosis, and modulating autophagy-related pathways ^28,29^. Mechanistically, BetA has been reported to disrupt mitochondrial function, promote reactive oxygen species (ROS) accumulation, and activate stress-responsive signaling cascades that contribute to cancer cell death ^30,31^. In addition, BetA has been shown to influence autophagy in a highly context-dependent manner. Although BetA has been reported to promote autophagy initiation by modulating AMP-activated protein kinase (AMPK), mechanistic target of rapamycin (mTOR), and Unc-51-like autophagy-activating kinase 1 (ULK1) ^32–35^, other studies suggest that BetA can disrupt autophagic flux, leading to dysfunctional or non-productive autophagy ^29,36^. These context-dependent effects suggest that autophagy regulation is an important component of BetA-mediated anticancer activity and may influence cancer cell responses to metabolic and therapeutic stress.

Despite these promising properties, the therapeutic application of BetA remains limited. BetA exhibits poor aqueous solubility, suboptimal pharmacokinetic properties, and limited target specificity, which collectively restrict its bioavailability and translational potential in clinical applications ^37^. These limitations have prompted increasing interest in the development of BetA-based derivatives and structurally optimized analogs aimed at enhancing potency, selectivity, and drug-like properties. To enhance antitumor activity, a variety of BetA–derived compounds have been developed, including C3-substituted betulin aldehydes ^38^, C-28-modified BetA ester derivatives ^39^, BetA analog 2c ^40^, SYK023 ^41^, and B5G1 ^42^.

In this study, we evaluated a series of previously reported BetA derivatives for their effects on UM cell viability and autophagy–lysosome function. We identified ALS-997 as a potent compound that impairs lysosome-dependent autophagic flux. Mechanistically, our findings suggest that ALS-997 engages Niemann–Pick disease type C1 (NPC1), perturbs lysosomal cholesterol homeostasis, and induces apoptotic signaling, thereby suppressing UM cell survival.

## Methods

### Cell culture

Human UM MP41 and primary epidermal melanocytes were purchased from American Type Culture Collection (Manassas, VA, USA), and 92.1 and Mel202 cells were from Sigma (St. Louis, MO, USA). MP41, 92.1, and Mel202 cells were cultured in RPMI 1640 (Corning, Glendale, AZ, USA) supplemented with 10% fetal bovine serum (FBS, Corning), and melanocytes were cultured in Dermal Cell Basal Medium supplemented with Adult Melanocyte Growth Kit and Antibiotic-Antimycotic (Thermo Fisher Scientific, Waltham, MA, USA) at 37 °C in a humidified incubator containing 5% CO_2_.

### Cell viability

The cells were seeded into a 96-well multiplate at a density of 1 × 10^4^ cells/well in the growth medium supplemented with 10% FBS 2 days before the experiment. Following treatment with agents for 24 h, cell viability was evaluated by Premix WST-1 Cell Proliferation Assay System (Takara Bio USA Inc., San Jose, CA, USA).

### Western blotting

The cells were washed twice with Phosphate-Buffered Saline (PBS), and the preparation of cell lysates was performed as described previously ^43^. Protein concentration in the cell extracts was determined using a bicinchoninic acid protein assay kit (Thermo Fisher Scientific). For Western blotting, proteins in the extract were separated by electrophoresis on a 10%, 12%, or 16% sodium dodecyl sulfate-polyacrylamide gel under reducing conditions and then electroblotted onto a polyvinylidene difluoride membrane. After blocking with PVDF Blocking Reagent for Can Get Signal (TOYOBO, Osaka, Japan) or 1% BSA, the membrane was allowed to react with the primary antibodies (1:1000). The immunoreactive proteins were visualized using peroxidase-conjugated secondary antibodies (1:5000) and SuperSignal West Pico PLUS (Thermo Fisher Scientific). The antibodies used in this study are described in Table S1. The band density was quantified with ImageJ software (National Institute of Health, Bethesda, MD, USA).

### Molecular modeling

Docking experiments were carried out using the MOE 2024.0601 (Chemical Computing Group ULC, Montreal, Canada). The cryo-EM structure of itraconazole-bound human NPC1 [PDB: 6UOX] was used as the receptor for the docking calculations. The BetA- and its derivatives-binding site of NPC1 was specified as being located at the interface between the middle luminal domain and the sterol-sensing domain. All atoms were protonated in an implicit solubilized environment at 300 K temperature, pH = 7, and a salt concentration of 0.1, using the Protonate3D module. The entire structure was energy minimized using the Amber10EHT force field, and BetA and its derivatives were docked to the binding site under the default settings, and fifty alternative conformations were constructed for BetA and its derivatives.

### Immunofluorescence staining

Immunofluorescence staining was performed as described previously ^44^. 92.1 cells were fixed with 4% phosphate-buffered paraformaldehyde solution for 10 min at room temperature and then permeabilized with PBS containing 0.1% Triton X-100 and 100 mM glycine for 10 min. Following treatment with PBS containing 0.1% Tween 20 and 1% BSA for 30 min, the cells were exposed to antibodies targeting LC3, p62, cleaved caspase-3, and LAMP2 (Table S1) for 24 h at 4°C, followed by incubation with donkey anti-rabbit IgG (H+L) highly cross-adsorbed secondary antibody conjugated with Alexa Fluor™ 488 and donkey anti-mouse IgG (H+L) highly cross-adsorbed secondary antibody conjugated with Alexa Fluor™ 594 (1:500, Thermo Fisher Scientific) for 1 h at room temperature. The coverslips were mounted in VECTASHIELD antifade mounting medium with DAPI (Vector Laboratories, Newark, CA, USA), and the immunolabeled and DAPI-stained cells were observed using LSM 880 and LSM 980 microscopes (Carl Zeiss, Oberkochen, Germany).

### Lysosomal staining with Lysotracker Green

Lysosomal staining was performed using Lysotracker Green DND-26 (Thermo Fisher Scientific). 92.1 cells were seeded on glass-bottom dishes and allowed to adhere overnight. After treatment with the indicated compounds for the specified time periods, cells were incubated with Lysotracker Green at a final concentration of 100 nM in pre-warmed culture medium for 30 min at 37 °C under 5% CO₂. Following incubation, cells were gently washed once with PBS and immediately imaged in fresh culture medium without fixation. Fluorescence images were acquired using a spinning disk live-cell imaging system (Carl Zeiss) under identical imaging conditions for all experimental groups. For quantitative analysis, Lysotracker fluorescence intensity was measured on a per-cell basis using ImageJ software. Individual cells were manually outlined as regions of interest (ROIs), and the mean fluorescence intensity within each ROI was calculated. Background fluorescence was determined from cell-free areas and subtracted from each measurement. At least 50 cells per condition were analyzed from three independent experiments.

### Cholesterol measurement

Filipin III (Cayman Chemical, Ann Arbor, MI, USA) was used to detect intracellular free cholesterol accumulation. Cells were washed with PBS and fixed with 4% paraformaldehyde for 15 min at room temperature. After fixation, cells were washed three times with PBS and incubated with Filipin III staining solution (50 μg/mL in PBS) for 1 h at room temperature in the dark. Following staining, cells were washed with PBS, and fluorescence signals were immediately observed using LSM980 confocal microscopes. Fluorescence intensity of Filipin III staining was quantified on a per-cell basis using ImageJ software. Confocal images were acquired under identical imaging conditions for all experimental groups. Individual cells were manually outlined as ROIs based on cell boundaries, and the mean fluorescence intensity of Filipin III within each ROI was measured. Background fluorescence was determined from cell-free regions and subtracted from each measurement. At least 50 cells were analyzed per condition from three independent experiments. Data are presented as mean fluorescence intensity per cell.

### Cellular thermal shift assay

A cellular thermal shift assay (CETSA) was performed to evaluate the interaction between ALS-997 and NPC1 in HEK293T cells. HEK293T cells, which express endogenous NPC1 at the physiological level, were seeded in 10 cm dishes, and cultured until they reached approximately 70–90% confluence on the day of the experiment. Cell lysates were treated with ALS-997 or DMSO for 2 h at 37°C. After treatment, cells were washed once with PBS and collected in PBS. Cell suspensions were centrifuged at 500 × g for 5 min at 4°C, and cell pellets were resuspended in PBS to obtain a homogeneous suspension. Equal volumes of the cell suspension were aliquoted into microcentrifuge tubes for each temperature point. The samples were heated for 3 min at the indicated temperatures using a thermal cycler, held at room temperature for 1 min, and then immediately transferred to ice. After thermal treatment, an equal volume of digitonin lysis buffer was added to each tube, and the samples were incubated on ice for 30 min with gentle mixing. Insoluble protein aggregates were removed by centrifugation at 20,000 × g for 20 min at 4°C. The soluble fractions were carefully collected without disturbing the pellets. NPC1 expression in the samples was evaluated by Western blot analysis. The band intensities were quantified using ImageJ software. The relative amount of soluble NPC1 at each temperature was normalized to the corresponding control protein or to the lowest temperature condition. Thermal stability curves were generated from three independent experiments.

### Quantitative real-time polymerase chain reaction (qPCR)

Total RNA was isolated from human dermal melanoma and the MP41, 92.1, and Mel202 cell lines using the RNeasy kit (QIAGEN, Hilden, Germany). Complementary DNA was synthesized using M-MLV reverse transcriptase (Thermo Fisher Scientific), and quantitative real-time PCR (qPCR) was performed using primers specific for human NPC1 and GAPDH.

### Statistical analysis

Data are presented as mean ± standard error (SE) from at least three independent experiments. Statistical analyses were performed using JMP software (JMP Student Edition 18.0, SAS Institute Inc., Cary, NC, USA). Statistical significance was determined using Student’s t-test (two-tailed) or one-way ANOVA followed by Tukey’s post hoc test. A p-value < 0.05 was considered statistically significant.

## Results

### Screening of cancer cell types which are sensitive for Bet A derivative treatment

First, we evaluated the anticancer activity of some of the BetA derivatives using the NCI-60 human tumor cell line panel. The compounds showed broad growth-inhibitory activity, with relatively consistent responses across cell lines derived from central nervous system (CNS) cancer, and melanoma (Fig. 1). In contrast, leukemia cell lines were generally less responsive. Responses varied substantially among individual cell lines derived from non-small cell lung, colon, ovarian, renal, prostate, and breast cancers (Fig. 1). Based on the relatively consistent response observed across the melanoma cell lines, we selected melanoma for further investigation. We focused specifically on uveal melanoma, a rare but life-threatening malignance that, unlike cutaneous melanoma, generally responds poorly to conventional chemotherapy and immunotherapy ^13^

**Figure 1.**
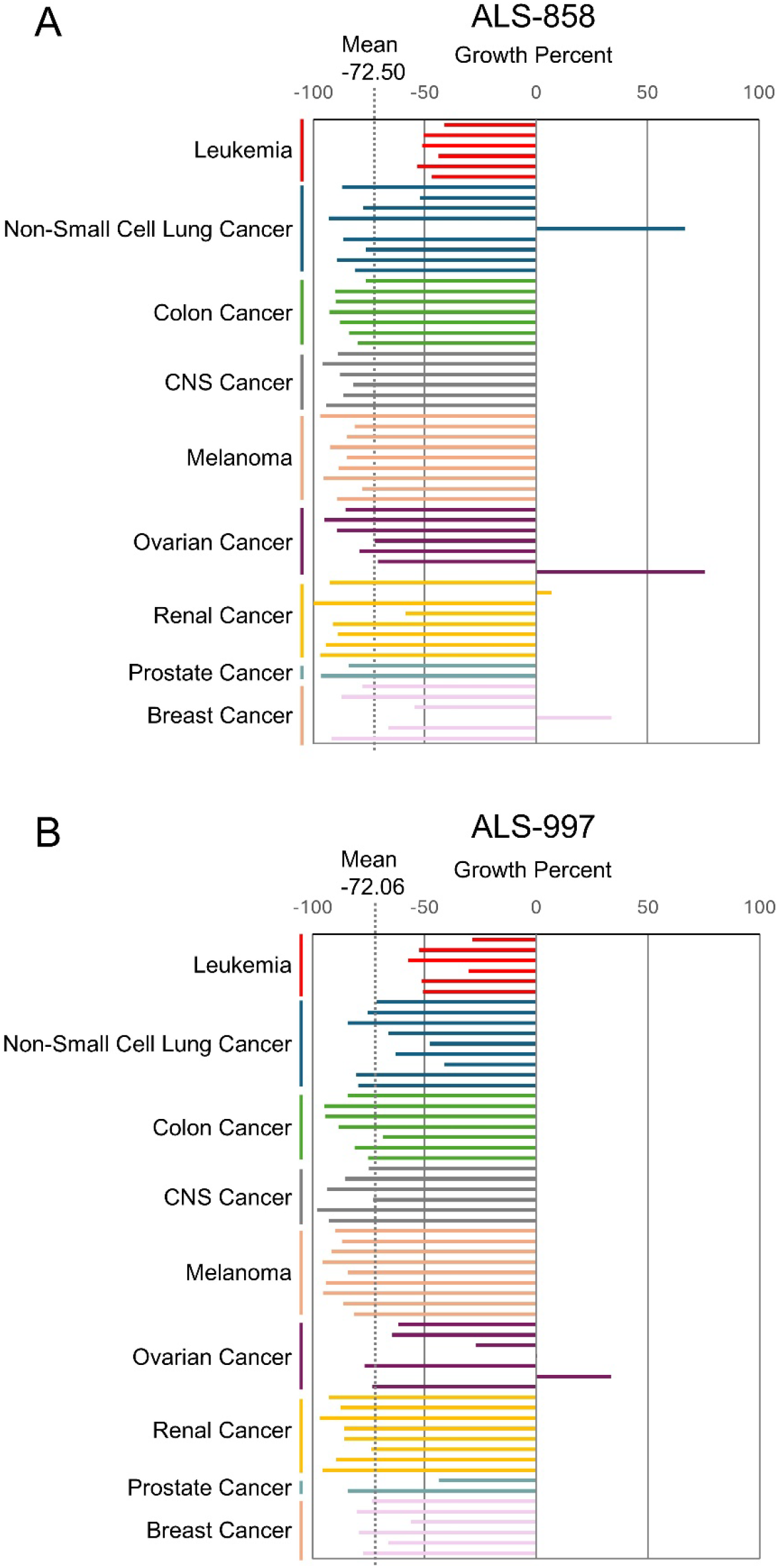
Growth-inhibitory profile of the BetA derivatives across the NCI-60 human tumor cell line panel. (A, B) Growth Percent values are shown for individual cell lines following treatment with the BetA derivatives, ALS-858 (A) and ALS-997 (B), at 10 µM for 48 hours. A Growth Percent value of 0 indicates no net growth relative to the cell population at the time of compound addition, whereas negative values indicate a reduction in cell number during the treatment period. The dotted gray line indicates the mean Growth Percent across the NCI-60 panel (−72.50% for ALS-858 and - 72.06% for ALS-997). Bars extending to the left of the mean line represent Growth Percent values below the panel mean, indicating a greater-than-average growth-inhibitory response to the compound. Data were obtained from the NCI-60 single-dose screen. The cell lines tested include leukemia (CCRF-CEM, HL-60(TB), K-562, MOLT-4, RPMI-8226, and SR), non-small cell lung cancer (A549, EKVX, HOP-62, HOP-92, NCI-H226, NCI-H23, NCI-H322M, NCI-H460, and NCI-H522), colon cancer (COLO 205, HCC-2998, NCT-116, NCT-15, NT29, KM12, and SW-620), CNS cancer (SF-268, SF-295, SF-539, SNB-19, SNB-75, and U251), cutaneous melanoma (LOX IMVI, MALME-3M, M14, MDA-MB-435, SK-MEL-2, SK-MEL-28, SK-MEL-5, UACC-257, and UACC-62), ovarian cancer (IGROV1, OVCAR-3, OVCAR-4, OVCAR-5, OVCAR-8, NCI/ADR-RES, and SK-OV-3), renal cancer (786-0, A498, ACHN, CAKI-1, RXF 393, SN12C, TK-10, and UO-31), prostate cancer (PC-3 and DU-145), and breast cancer (MCF7, MDA-MB-231, HS 578T, BT-549, T-47D, and MDA-MB-468).

### Betulinic acid impairs autophagic flux by disrupting lysosomal function

Previous studies have reported seemingly divergent effects of BetA on autophagy, with evidence supporting both enhanced autophagy initiation and autophagy flux inhibition ^29,32–36^. To clarify its role, we first examined the effects of BetA on autophagy in UM 92.1 cells. BetA treatment significantly reduced cell viability at concentrations of 50 μM or higher (Fig. 2A). p62 is an autophagy adaptor protein that binds ubiquitinated protein aggregates and damaged organelles, delivering them to the autophagosome membrane, where they are selectively degraded through interaction with LC3 ^45^. BetA treatment markedly increased p62 protein levels and elevated the LC3-II/LC3-I ratio at concentrations of 20 μM or higher (Fig. 2B, C).

**Figure 2.**
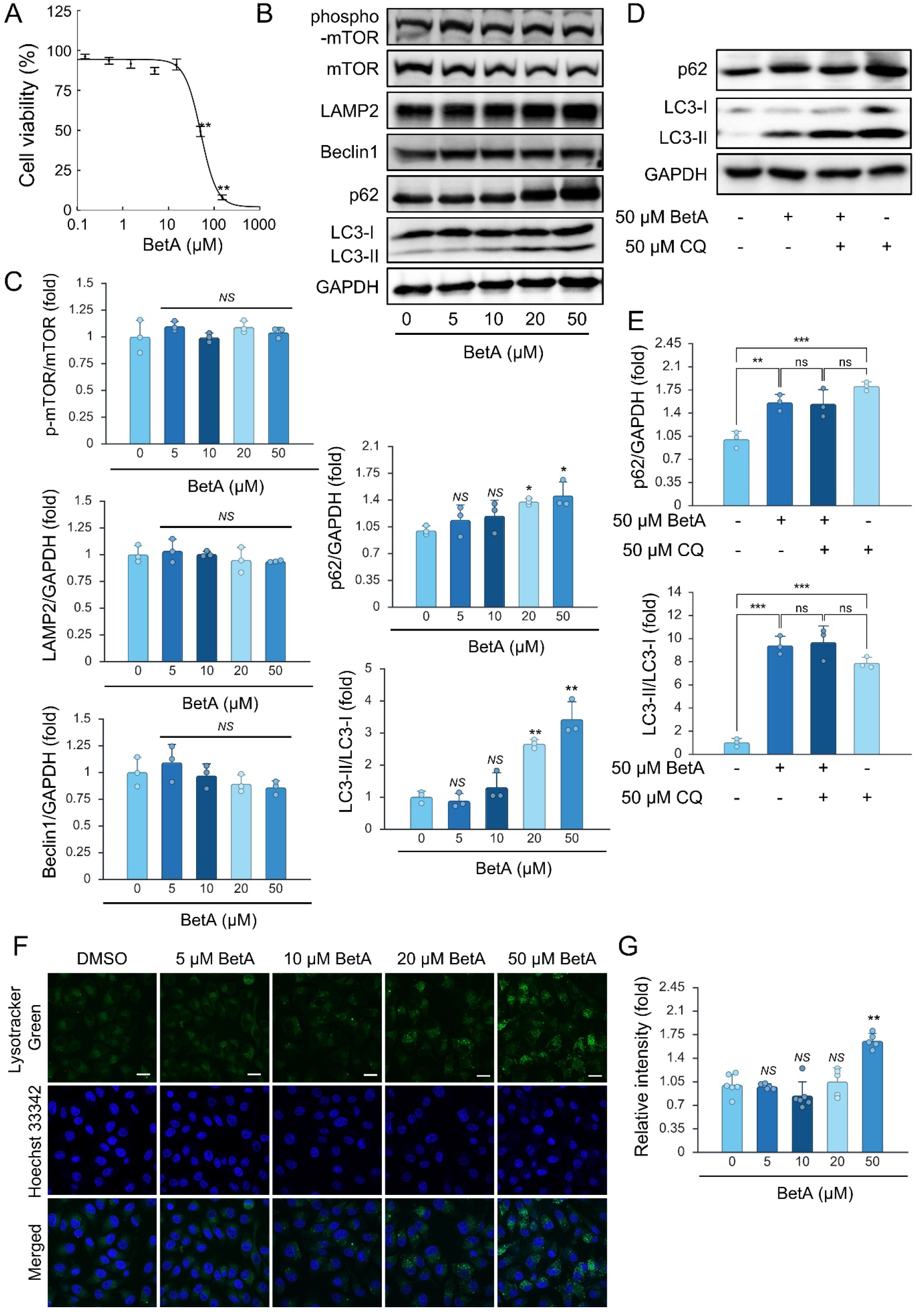
Betulinic acid impairs autophagic flux by disrupting lysosomal function. (A) WST-1 assay showing that BetA reduces the viability of UM 92.1 cells in a dose-dependent manner. n=8 (B, C) Western blot analysis showing that BetA dose-dependently increases the LC3-II/LC3-I ratio and p62 protein levels, whereas mTOR phosphorylation, LAMP2 expression, and Beclin1 expression are not significantly affected. n=3. Quantification of phosphorylated mTOR (p-mTOR) relative to total mTOR, p62, LAMP2, and Beclin1 normalized to GAPDH, and the LC3-II/LC3-I ratio is presented as mean ± SE. (D, E) Western blot analysis showing the effect of chloroquine (CQ) co-treatment with BetA. Co-treatment with BetA and CQ did not further increase the LC3-II/LC3-I ratio compared to CQ alone. n=3. (F, G) Lysotracker Green staining of lysosomes and Hoechst 33342 staining of nuclei in 92.1 cells treated with 0-50 µM of BetA. Scale bar = 20 µm. Lysotracker Green fluorescence intensity was quantified and is presented as mean ± SE. n=6. Statistical significance was determined by one-way ANOVA. * p<0.05, ** p<0.01, *** p<0.001, and NS, not significant.

In contrast, BetA did not significantly alter mTOR phosphorylation, as indicated by the p-mTOR/mTOR ratio, nor did it affect the expression of LAMP2 or Beclin1 (Fig. 2B, C). Chloroquine (CQ), which increases lysosomal pH and blocks autophagosome degradation, is commonly used to assess autophagic flux by promoting LC3-II accumulation ^46^. Co-treatment with BetA and CQ did not further increase the LC3-II/LC3-I ratio compared to CQ alone (Fig. 2D, E), indicating that BetA does not enhance autophagic flux. Moreover, lysosomal staining with Lysotracker Green revealed a significant increase in fluorescence intensity following treatment with 50 μM BetA, accompanied by enlarged and clustered lysosomal structures (Fig. 2F). Collectively, these findings suggest that in UM cells, BetA impairs autophagic flux by impairing lysosomal function rather than through modulation of mTOR signaling.

### Evaluation of BetA derivatives and their antitumor activity in UM cells

To improve the physicochemical properties, antitumor activity, and tumor selectivity of BetA, a series of BetA derivatives were designed and synthesized as described (Fig. 3) ^47–49^. Compared to the parent compound BetA, all synthesized BetA derivatives, except for ALS-769, exhibited reduced IC₅₀ values against the UM cell lines MP41, 92.1, and Mel202 (Table 1). Notably, among these derivatives, ALS-997 and ALS-1022 showed preferential cytotoxicity toward selected UM cell lines relative to primary epidermal melanocytes. Moreover, the physicochemical properties of the ALS compounds were calculated using SwissADME (https://www.swissadme.ch/) and ADMETlab2.0 (https://admetmesh.scbdd.com/) and compared to those of betulinic acid (BetA) (Table 2). BetA showed a SlogP value of 7.09. SlogP is a predicted measure of lipophilicity based on octanol–water partitioning and can influence aqueous solubility, membrane partitioning, and intracellular distribution. Most ALS compounds exhibited SlogP values comparable to BetA (6.5–7.5), suggesting that they largely retain BetA-like hydrophobic properties. Higher lipophilicity was observed for ALS-858, ALS-1025, and ALS-1117 (SlogP > 8.5), which may favor stronger association with lipid-rich membranes but could also reduce aqueous solubility. In contrast, ALS-987 and ALS-1021 displayed lower SlogP values (≤5.5), indicating relatively improved aqueous solubility but reduced lipid membrane association.

**Figure 3.**
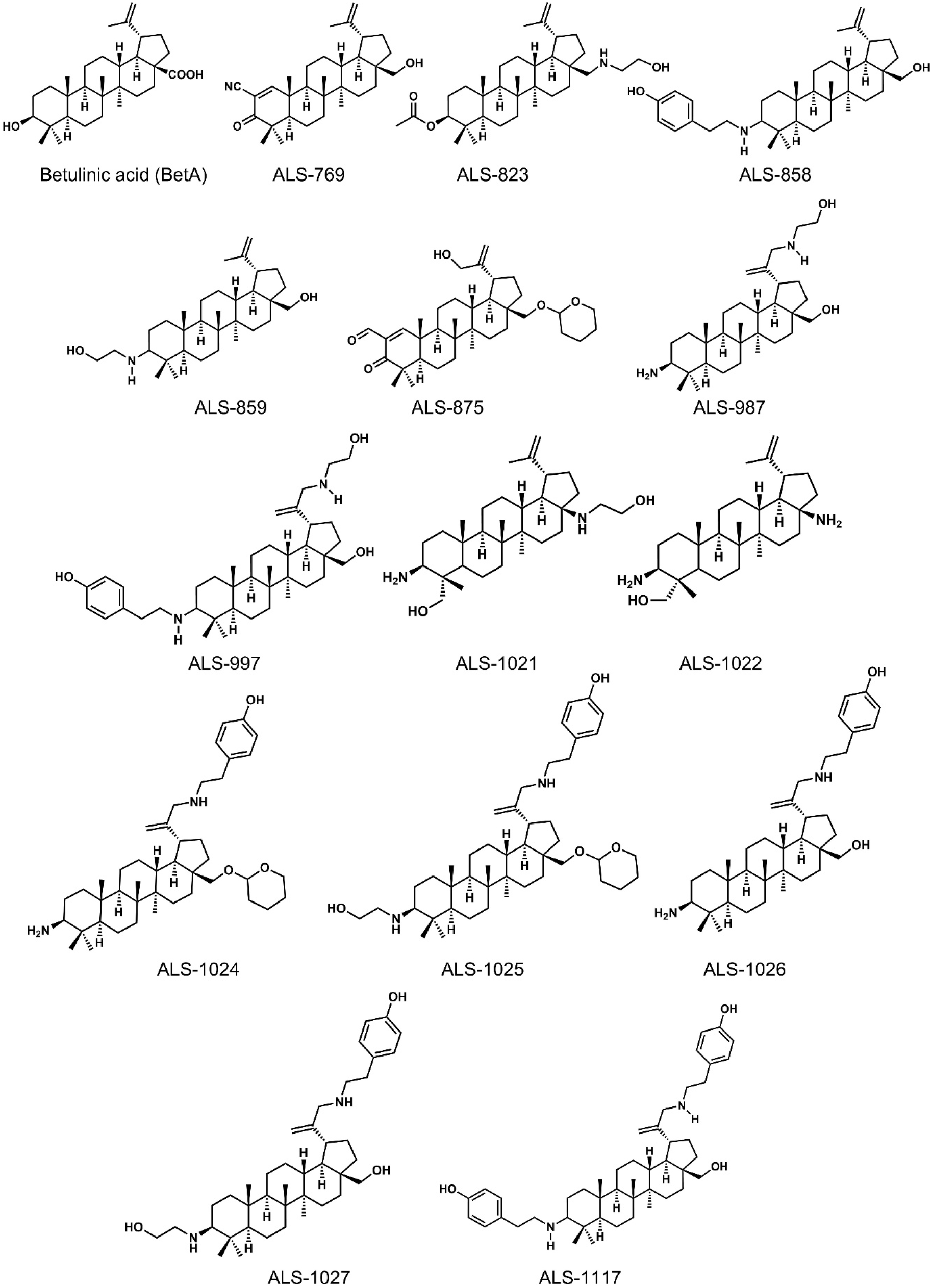
Structures of BetA derivatives. Betulinic acid (BetA, Mw = 456.71 g/mol), ALS-769 (Mw = 465.7), ALS-823 (Mw = 527.7), ALS-858 (Mw = 561.9), ALS-859 (Mw = 485.7), ALS-875 (Mw = 566.81), ALS-987 (Mw = 500.8), ALS-997 (Mw = 620.95), ALS-1021 (Mw = 486.7), ALS-1022 (Mw = 442.7), ALS-1024 (Mw = 661.0), ALS-1025 (Mw = 705.0), ALS-1026 (Mw = 576.9), ALS-1027 (Mw = 621.0), and ALS-1117 (Mw = 697.05).

**Table 1.**
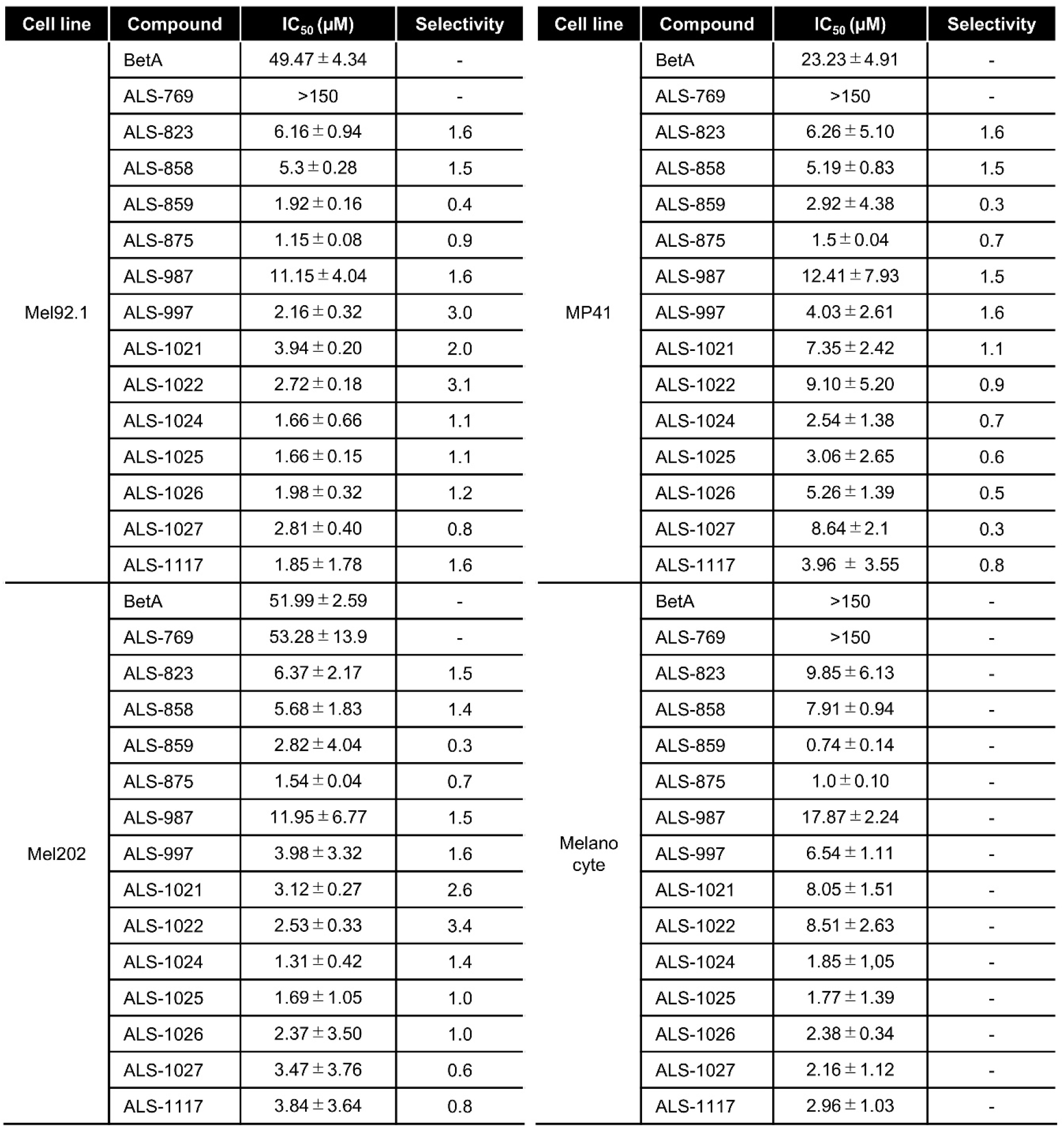
IC₅₀ values of BetA-derivatives in UM cells and human melanocytes. Three uveal melanoma cell lines and human melanocytes were treated with increasing concentrations of each BetA derivative for 24 h, followed by assessment of cell viability using the WST-1 assay. IC₅₀ values were calculated from dose–response curves and are shown as mean ± SE from independent experiments. N=8. Selectivity indices were calculated by dividing the IC₅₀ value in human melanocytes by the IC₅₀ value in each uveal melanoma cell line.

| Cell line | Compound | IC<sub>50</sub> (µM) | Selectivity | Cell line | Compound | IC<sub>50</sub> (µM) | Selectivity |
|---|---|---|---|---|---|---|---|
| Mel92.1 | BetA | 49.47 ± 4.34 | - | MP41 | BetA | 23.23 ± 4.91 | - |
|  | ALS-769 | >150 | - |  | ALS-769 | >150 | - |
|  | ALS-823 | 6.16 ± 0.94 | 1.6 |  | ALS-823 | 6.26 ± 5.10 | 1.6 |
|  | ALS-858 | 5.3 ± 0.28 | 1.5 |  | ALS-858 | 5.19 ± 0.83 | 1.5 |
|  | ALS-859 | 1.92 ± 0.16 | 0.4 |  | ALS-859 | 2.92 ± 4.38 | 0.3 |
|  | ALS-875 | 1.15 ± 0.08 | 0.9 |  | ALS-875 | 1.5 ± 0.04 | 0.7 |
|  | ALS-987 | 11.15 ± 4.04 | 1.6 |  | ALS-987 | 12.41 ± 7.93 | 1.5 |
|  | ALS-997 | 2.16 ± 0.32 | 3.0 |  | ALS-997 | 4.03 ± 2.61 | 1.6 |
|  | ALS-1021 | 3.94 ± 0.20 | 2.0 |  | ALS-1021 | 7.35 ± 2.42 | 1.1 |
|  | ALS-1022 | 2.72 ± 0.18 | 3.1 |  | ALS-1022 | 9.10 ± 5.20 | 0.9 |
|  | ALS-1024 | 1.66 ± 0.66 | 1.1 |  | ALS-1024 | 2.54 ± 1.38 | 0.7 |
|  | ALS-1025 | 1.66 ± 0.15 | 1.1 |  | ALS-1025 | 3.06 ± 2.65 | 0.6 |
|  | ALS-1026 | 1.98 ± 0.32 | 1.2 |  | ALS-1026 | 5.26 ± 1.39 | 0.5 |
|  | ALS-1027 | 2.81 ± 0.40 | 0.8 |  | ALS-1027 | 8.64 ± 2.1 | 0.3 |
|  | ALS-1117 | 1.85 ± 1.78 | 1.6 |  | ALS-1117 | 3.96 ± 3.55 | 0.8 |
| Mel202 | BetA | 51.99 ± 2.59 | - | Melanocyte | BetA | >150 | - |
|  | ALS-769 | 53.28 ± 13.9 | - |  | ALS-769 | >150 | - |
|  | ALS-823 | 6.37 ± 2.17 | 1.5 |  | ALS-823 | 9.85 ± 6.13 | - |
|  | ALS-858 | 5.68 ± 1.83 | 1.4 |  | ALS-858 | 7.91 ± 0.94 | - |
|  | ALS-859 | 2.82 ± 4.04 | 0.3 |  | ALS-859 | 0.74 ± 0.14 | - |
|  | ALS-875 | 1.54 ± 0.04 | 0.7 |  | ALS-875 | 1.0 ± 0.10 | - |
|  | ALS-987 | 11.95 ± 6.77 | 1.5 |  | ALS-987 | 17.87 ± 2.24 | - |
|  | ALS-997 | 3.98 ± 3.32 | 1.6 |  | ALS-997 | 6.54 ± 1.11 | - |
|  | ALS-1021 | 3.12 ± 0.27 | 2.6 |  | ALS-1021 | 8.05 ± 1.51 | - |
|  | ALS-1022 | 2.53 ± 0.33 | 3.4 |  | ALS-1022 | 8.51 ± 2.63 | - |
|  | ALS-1024 | 1.31 ± 0.42 | 1.4 |  | ALS-1024 | 1.85 ± 1,05 | - |
|  | ALS-1025 | 1.69 ± 1.05 | 1.0 |  | ALS-1025 | 1.77 ± 1.39 | - |
|  | ALS-1026 | 2.37 ± 3.50 | 1.0 |  | ALS-1026 | 2.38 ± 0.34 | - |
|  | ALS-1027 | 3.47 ± 3.76 | 0.6 |  | ALS-1027 | 2.16 ± 1.12 | - |
|  | ALS-1117 | 3.84 ± 3.64 | 0.8 |  | ALS-1117 | 2.96 ± 1.03 | - |

**Table 2:** The physicochemical properties of the ALS compounds and BetA, calculated using SwissADME and ADMETlab2.0. (separate Excel file)

BetA showed a total polar surface area (tPSA) of 57.53. Compared to BetA, the ALS compounds generally exhibited greater polarity, with tPSA values exceeding 70 for most analogs and reaching approximately 85 for several. These changes were accompanied by increased hydrogen-bonding capacity, with ALS compounds typically possessing 3–6 hydrogen-bond acceptors and 1–5 hydrogen-bond donors, compared to 2 acceptors and 3 donors in BetA. These results suggest that structural modification of BetA yielded ALS compounds with physicochemical properties more consistent with a drug-like profile than those of the parent compound.

Next, the effect of BetA-derived compounds on autophagic flux was evaluated based on the LC3-II/LC3-I ratio. Among the ALS compounds tested, 3 µM of ALS-859, ALS-997, ALS-1024, and ALS-1025 exhibited markedly increased LC3-II/LC3-I ratios, indicating pronounced inhibition of autophagic flux (Fig. 4A, B). In contrast, parental BetA showed little to no effect at the same concentration. Consistent with these findings, treatment with ALS-859, ALS-997, ALS-1024, ALS-1025, ALS-1026, ALS-1027, and ALS-1117 resulted in a significant increase in Lysotracker fluorescence intensity, accompanied by prominent lysosomal clustering, as observed by fluorescence microscopy (Fig. 4C, D).

**Figure 4.**
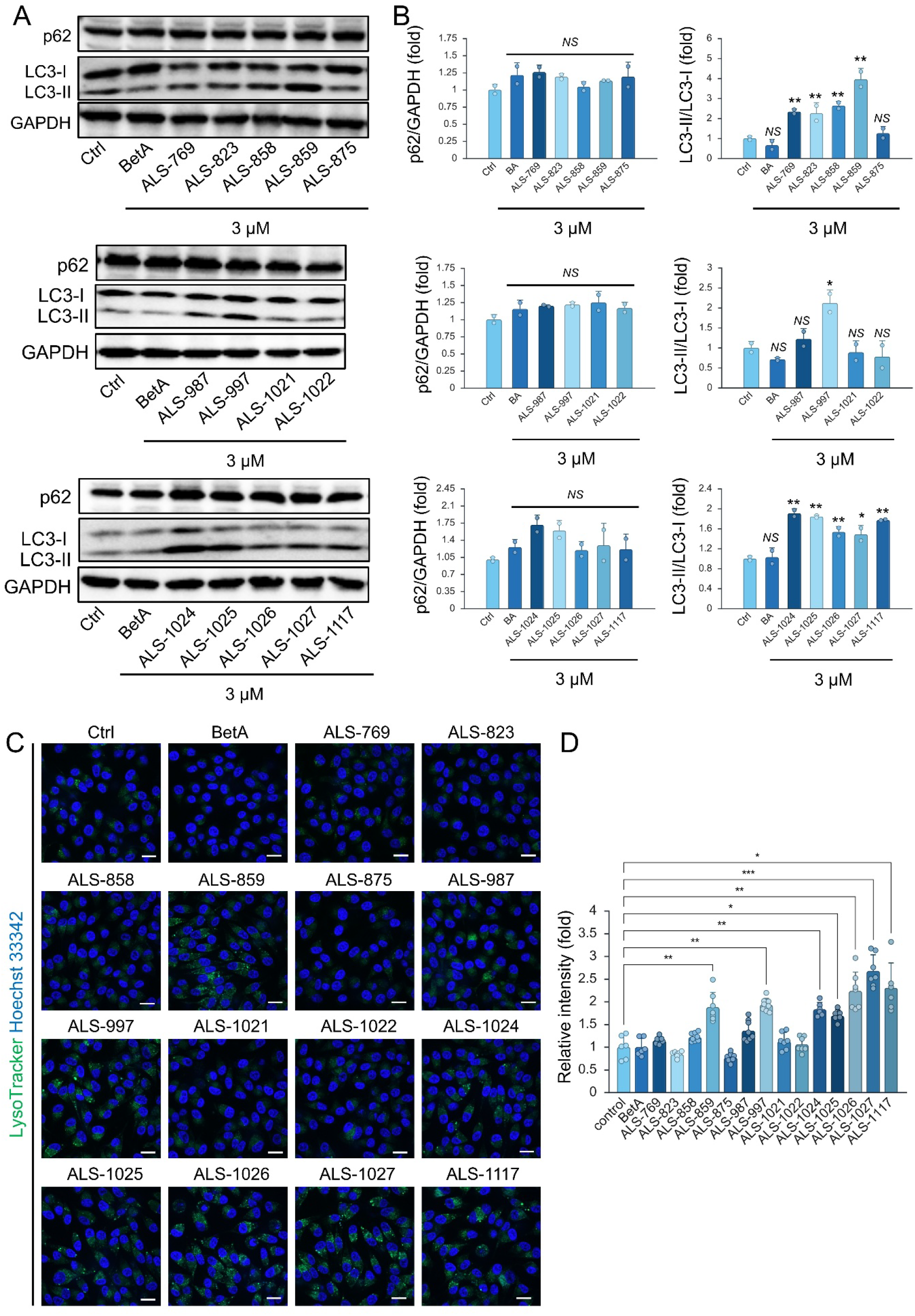
Screening of BetA derivatives based on autophagy- and lysosome-related markers. (A, B) Western blot analysis showing that some BetA derivatives increase the LC3-II/LC3-I ratio at low concentration (3 µM), at which concentration BetA does not show any increase. P62 increase was not observed at this concentration. Quantification of p62 normalized to GAPDH, and the LC3-II/LC3-I ratio is presented as mean ± SE. n=2. (C, D) Lysotracker Green staining of lysosomes and Hoechst 33342 staining of nuclei in 92.1 cells treated with 2 µM of BetA derivatives for 12 h. Scale bar = 20 µm. Lysotracker Green fluorescence intensity was quantified and is presented as mean ± SE. n=6-8. Statistical significance was determined by one-way ANOVA. * p<0.05, ** p<0.01, *** p<0.001, and NS, not significant.

### ALS-997 activates apoptotic signaling and induces cell death

Based on these findings, ALS-997 was selected for further mechanistic studies owing to its potent inhibitory activity on autophagy and preferential activity toward 92.1 cells relative to primary epidermal melanocyte. ALS-997 significantly reduced cell viability in 92.1 cells at concentrations ≥1.5 µM. In contrast, a significant decrease in viability was observed only at concentrations ≥15 µM in normal melanocytes (Fig. 5A). Next, the effects of BetA and ALS-997 on apoptosis-related signaling were evaluated in 92.1 cells. Treatment with ALS-997 at 5 µM resulted in significant increases in the cleaved PARP/PARP and cleaved caspase-3 immunofluorescence, whereas BetA at the same concentration did not induce significant changes in these markers (Fig. 5B, C). Collectively, these results indicate that ALS-997 activates apoptotic signaling and induces apoptosis at concentrations lower than those of BetA.

**Figure 5.**
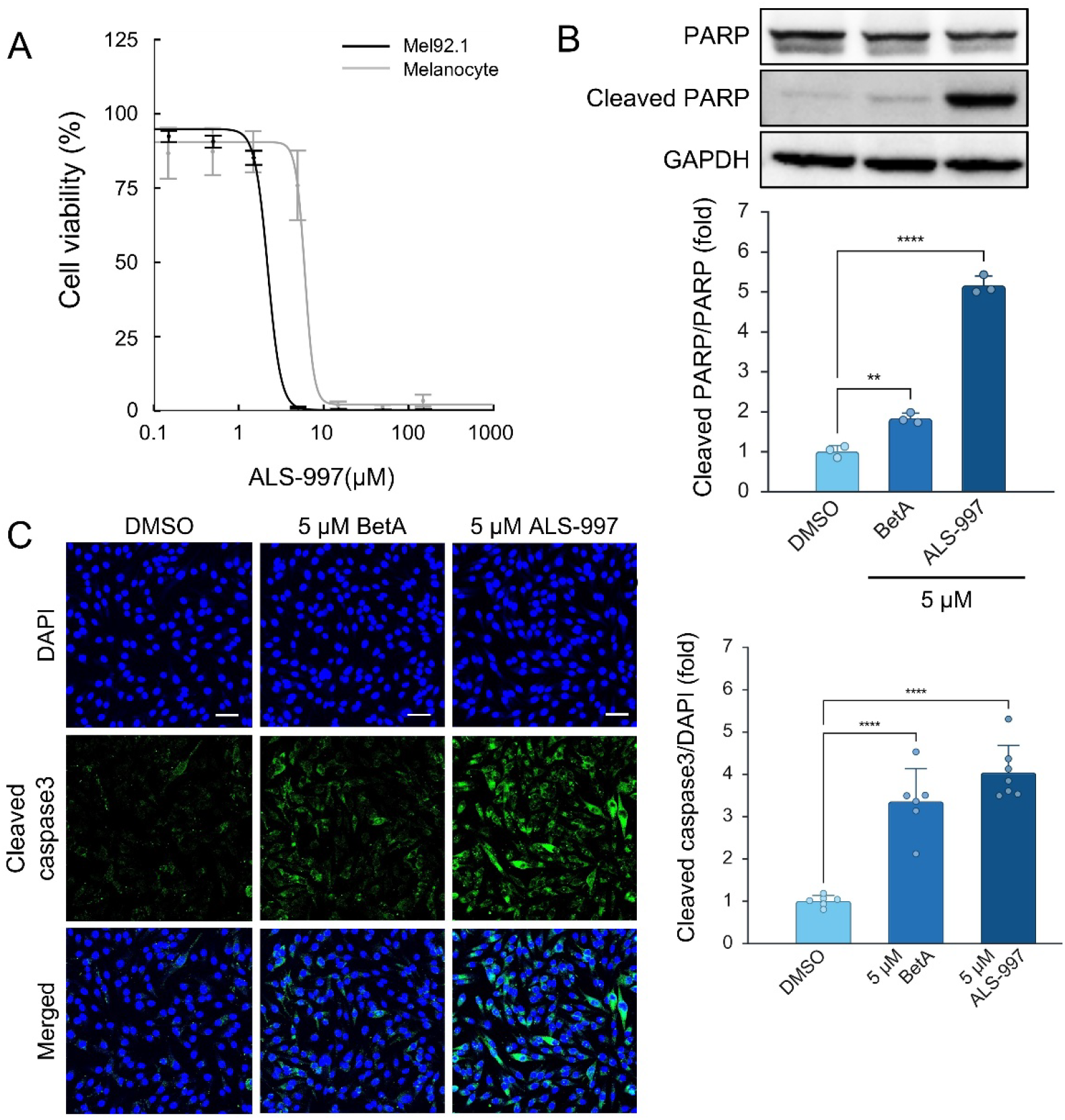
ALS-997 activates apoptotic signaling and induces cell death. (A) WST-1 assay showing that ALS-997 selectively reduces the viability of UM 92.1 cells compared to normal human melanocytes. n=8. (B) Western blot analysis showing that 5 µM of BetA modestly increases PARP cleavage, whereas 5 µM of ALS-997 markedly induces PARP cleavage. n=3. (C) Immunostaining of cleaved caspase-3 showing that 5 µM of BetA modestly increases caspase-3 cleavage, whereas 5 µM of ALS-997 markedly induces caspase-3 cleavage. n=5-7. One-way ANOVA. ** p<0.01, *** p<0.001, and **** p<0.0001.

### NPC1 inhibition by ALS-997 causes lysosomal impairment and autophagy blockade

Finally, to further investigate how ALS-997 impairs autophagic flux, we evaluated its effects on lysosomal function. ALS-997 induced a concentration-dependent increase in p62 levels and significantly elevated the LC3-II/LC3-I ratio at concentrations ≥2 µM (Fig. 6A, B). In parallel, ALS-997 significantly reduced LAMP2 expression, a lysosomal marker. In contrast, no significant changes in the p-mTOR/mTOR ratio were observed, consistent with the effects of BetA. Co-treatment with ALS-997 and CQ did not further increase the LC3-II/LC3-I ratio compared to CQ treatment alone (Fig. 6C, D). LC3 and p62 puncta accumulation was also observed in cells treated with ALS-997 in a dose-dependent manner (Fig. 6E, F). Moreover, Lysotracker staining revealed that ALS-997 increased lysosomal fluorescence intensity and induced lysosomal clustering at concentrations up to 2 µM, whereas lysosomal fluorescence was almost completely abolished at 5 µM (Fig. 6G, H). Collectively, these results suggest that ALS-997 impairs autophagic flux by disrupting lysosomal function.

**Figure 6.**
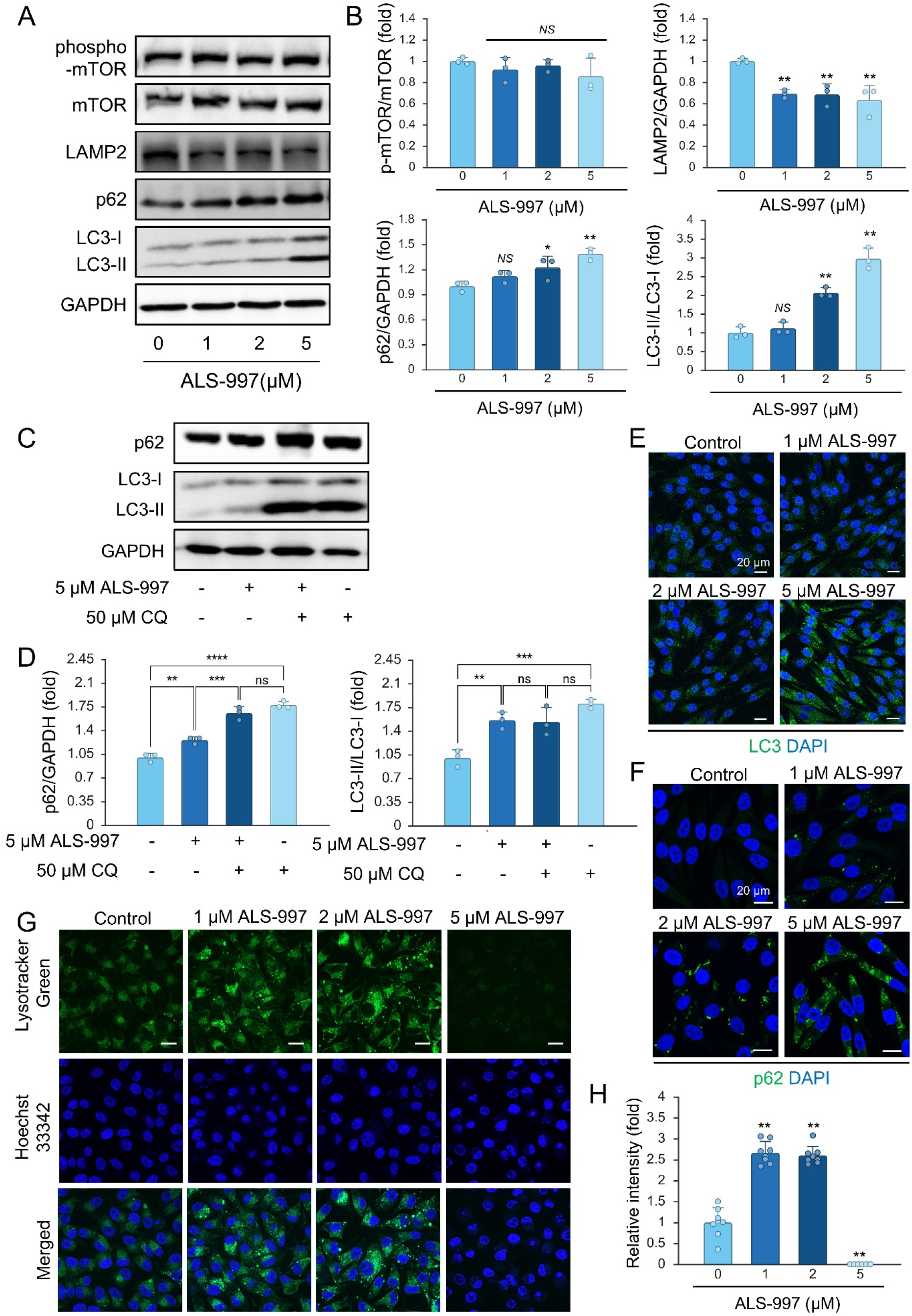
ALS-997 inhibits autophagic flux by blocking lysosomal function. (A, B) Western blot analysis of 92.1 lysate showing that ALS-997 dose dependently increases LC3-II/LC3-I ratio and p62 expression, whereas LAMP2 expression is reduced by 1 µM and remains similarly reduced by 2 and 5 µM. Quantification of phosphorylated mTOR relative to total mTOR, p62, and LAMP2 normalized to GAPDH, and the LC3-II/LC3-I ratio is presented as mean ± SE. n=3. (C, D) Western blot analysis showing the effect of CQ co-treatment with ALS-997. Co-treatment with ALS-997 and CQ did not further increase the LC3-II/LC3-I ratio compared to CQ alone. n=3. (E, F) Immunostaining of LC3 or p62 together with DAPI showing that ALS-997 dose-dependently increases LC3 and p62 in 92.1 cells. (G, H) Lysotracker Green staining of lysosomes and Hoechst 33342 staining of nuclei in 92.1 cells treated with 0-5 µM of ALS-997. Scale bar = 20 µm. Lysotracker Green fluorescence intensity was quantified and is presented as mean ± SE. n=6. Statistical significance was determined by one-way ANOVA. * p<0.05, ** p<0.01, *** p<0.001, **** p<0.0001, and NS, not significant.

Notably, similar lysosomal phenotypes—characterized by perinuclear accumulation and clustering of enlarged lysosomes, often accompanied by increased Lysotracker fluorescence intensity—have been reported upon pharmacological or genetic inhibition of NPC1 ^50,51^. NPC1 is a cholesterol transporter in the lysosome ^52^. Accordingly, cholesterol accumulation induced by ALS-997 was evaluated by Filipin staining. In Mel92.1 cells, treatment with ALS-997 markedly increased Filipin III fluorescence intensity, comparable to that observed upon treatment with U18666A, a well-established NPC1 inhibitor, suggesting intracellular cholesterol accumulation. (Fig. 7A, B). Next, to investigate whether ALS-997 inhibits NPC1 through direct binding, a CETSA was performed. CETSA is a target engagement assay that directly evaluates compound–protein interactions in cells by measuring changes in protein thermal stability; binding of a compound to its target protein typically results in increased thermal stability. In DMSO-treated cells, soluble NPC1 levels decreased with increasing temperature. In contrast, treatment with ALS-997 and U18666A increased the amount of soluble NPC1 remaining at higher temperatures. Notably, NPC1 signals were maintained at higher levels in ALS-997- and U18666A-treated cells than in DMSO-treated cells around 57.3 – 59.7 °C. These results indicate that ALS-997 thermally stabilizes NPC1 in cells, suggesting that ALS-997 directly or closely interacts with NPC1. (Fig. 7C, D). To elucidate the structural determinants underlying NPC1 inhibition by ALS-997, we constructed docking models of cholesterol, U18666A, and ALS-997 bound to NPC1 (Fig. 8 and Fig. S1-2). ALS-997 is positioned near the interface between the middle luminal domain and the sterol-sensing domain, overlapping with the cholesterol-binding site (Fig. 8A, B), similar to U18666A (Fig. S2). Furthermore, molecular docking analysis suggested that the hydroxymethyl group at the C13 position of ALS-997 forms a hydrogen bond with Glu688 of NPC1. In contrast, the phenolic hydroxyl group engages in hydrogen bonding with Ile609, thereby contributing to its inhibitory activity (Fig. 8C). The calculated docking scores for cholesterol, BetA, and ALS-997 were -7.59, -6.81, and -8.28, respectively, indicating a stronger predicted binding affinity of ALS-997 to NPC1 compared to cholesterol and BetA. Collectively, these findings suggest that ALS-997 inhibits NPC1, leading to lysosomal cholesterol accumulation and subsequent lysosomal dysfunction, which in turn results in inhibition of autophagic flux.

**Figure 7.**
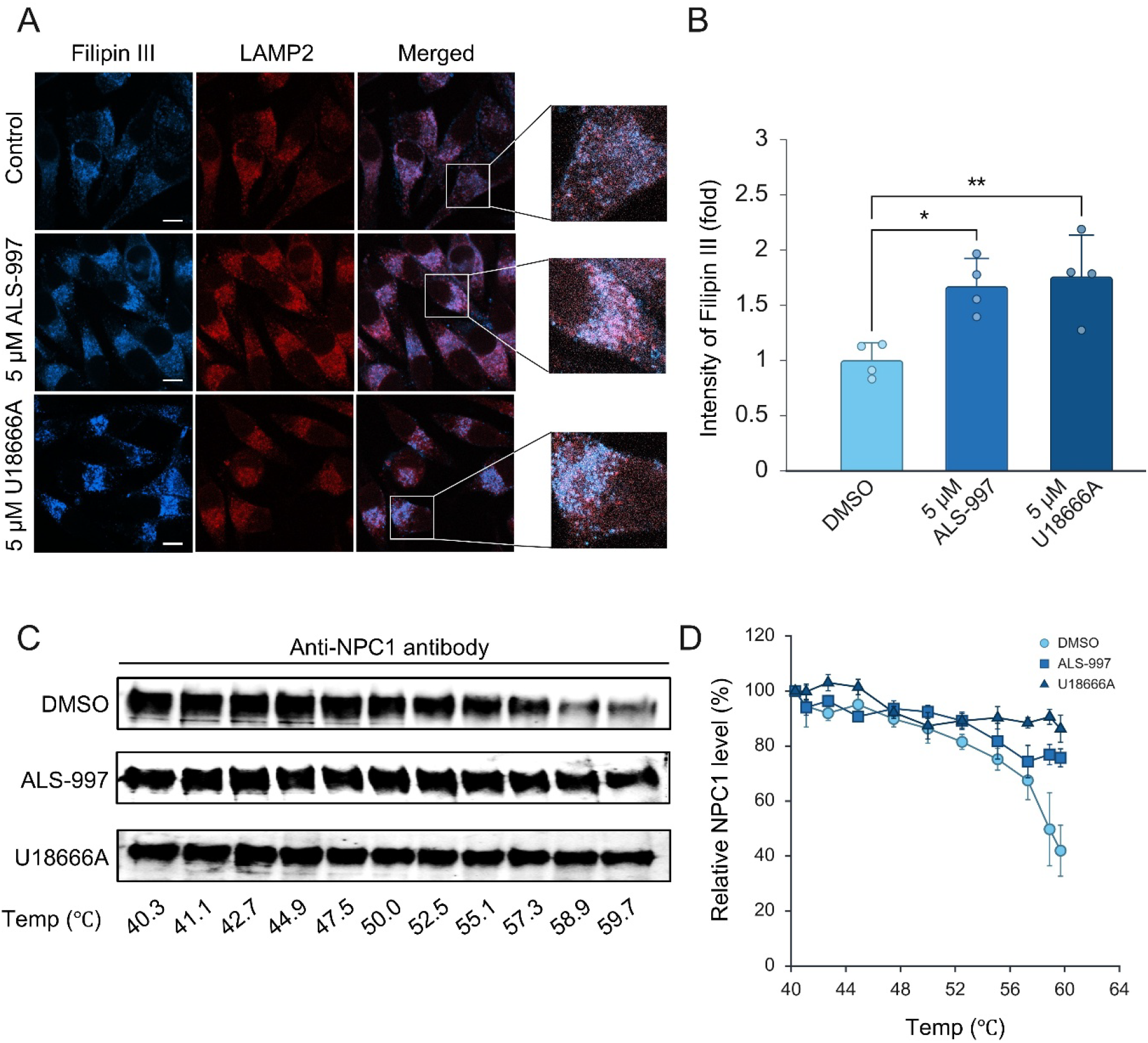
The inhibition of NPC1 by ALS-997. (A, B) Filipin III (cholesterol) and LAMP2 (lysosome) staining showing that 5 µM of ALS-997 induces cholesterol accumulation in lysosomes in 92.1 cells. A positive control U18666A, an inhibitor for lysosomal cholesterol transporter NPC1, also increased cholesterol accumulation in lysosomes. Scale bar = 10 µm. Fold increase of Filipin III intensity is shown as mean ± SE. n=4. (C, D) Thermal shift assay showing that ALS-997 interacts with NPC1. HEK293T cell lysates expressing NPC1 were incubated with DMSO, ALS-997, or U18666A (positive control), followed by heating at the indicated temperatures. NPC1 levels remaining after thermal challenge were analyzed by western blotting and quantified. ALS-997, similar to U18666A, increased the thermal stability of NPC1, suggesting direct or indirect interaction with NPC1. Data are shown as mean ± SE. n=3.

**Figure 8.**
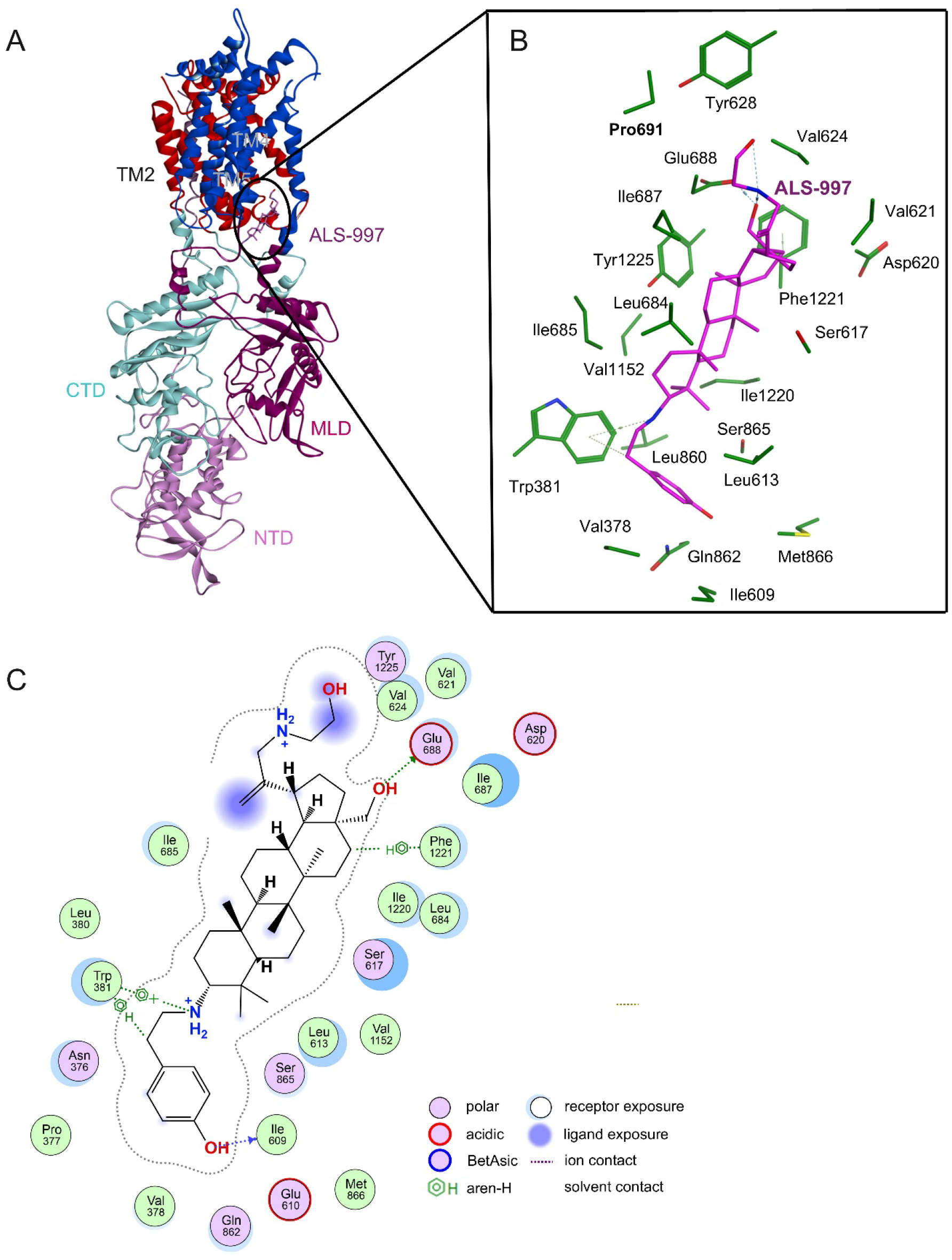
Docking model of NPC1 bound to ALS-997. (A) Overall docking model of ALS-997 bound to human NPC1 structure [PDB: 6UOX]. NPC1 domains are shown in different colors: transmembrane helices/domain (TM), N-terminal luminal domain (NTD), middle luminal domain (MLD), and C-terminal luminal domain (CTD). ALS-997 is shown in magenta. (B) Enlarged view of the predicted ALS-997 binding pocket located near the interface between the MLD and the sterol-sensing domain-containing transmembrane region. Amino acid residues surrounding ALS-997 are shown as sticks. (C) Two-dimensional ligand-protein interaction diagram of ALS-997 with NPC1. The docking model predicts hydrogen-bond interactions between ALS-997 and NPC1 residues, including Glu688 and Ile609. Docking calculations were performed using MOE2024.0601.

Interestingly, NPC1 expression was elevated in UM cell lines compared with normal melanocytes at both the mRNA and protein levels (Fig. 9A, B). This differential expression may contribute, at least in part, to the preferential activity of ALS-997 against UM cells and further supports NPC1 as a functionally relevant target of the compound. Taken together, our findings demonstrate that pharmacological inhibition of NPC1 impairs lysosome-dependent autophagic flux, thereby suppressing UM cell survival, and support NPC1 as a potential therapeutic target in UM.

**Figure 9.**
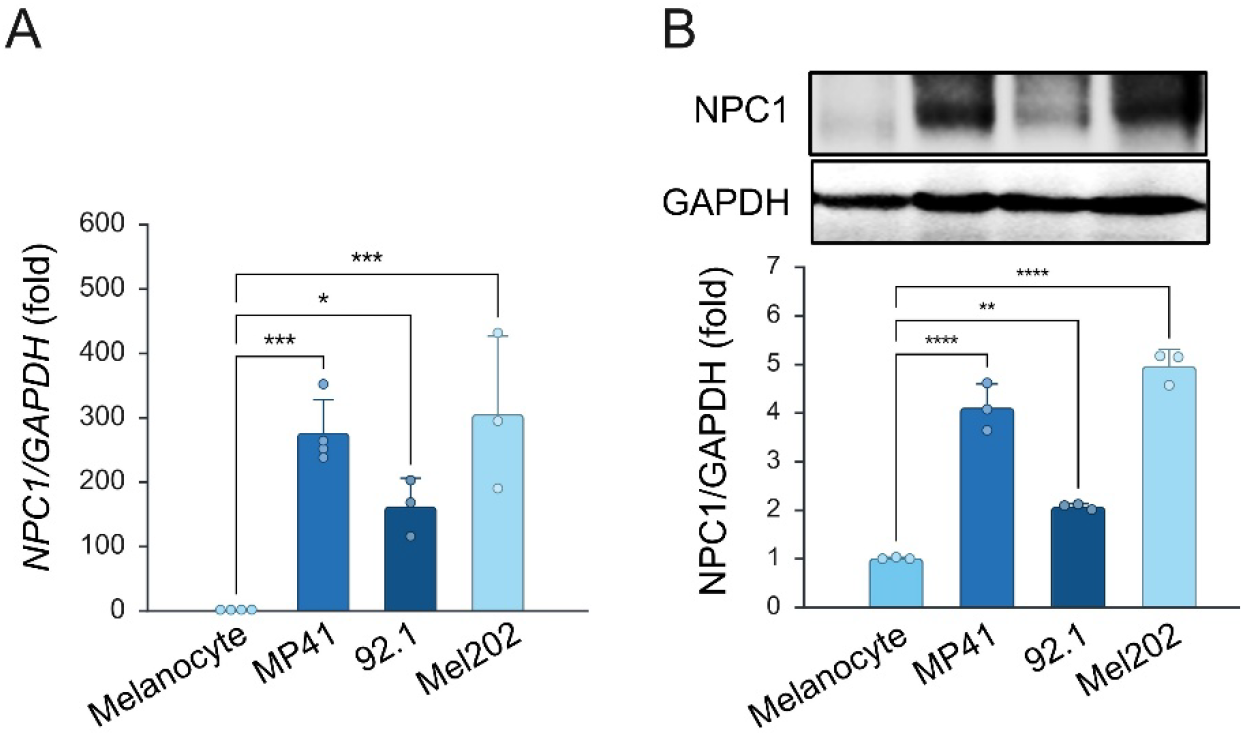
NPC1 is upregulated in UM. (A) NPC1 mRNA expression was measured by qPCR in UM cell lines (MP41, 92.1, and Mel202) and normal melanocytes. Expression levels were normalized to GAPDH. Data are shown as mean ± SE; n=3-4. (B) Representative western blot analysis of NPC1 protein expression in UM cell lines compared with normal melanocytes. Quantification was performed from three independent experiments, and shown as mean ± SE; n=3. One-way ANOVA. * p<0.05, ** p<0.01, *** p<0.001, and **** p<0.0001.

## Discussion

Autophagy plays a context-dependent and often paradoxical role in cancer, acting as both a tumor-suppressive and tumor-promoting mechanism depending on tumor stage, genetic background, and microenvironmental stress ^53^. In established cancers, including UM, autophagy frequently functions as a cytoprotective stress-adaptation pathway that supports tumor cell survival, metabolic flexibility, and therapeutic resistance ^23,54^. This dual nature of autophagy has complicated the development of autophagy-targeting anticancer strategies and underscores the need for precise modulation of autophagic flux rather than indiscriminate activation or inhibition. Consistent with this concept, accumulating evidence suggests that pharmacological inhibition of autophagy can sensitize cancer cells to metabolic and therapeutic stress, particularly in tumors with high autophagy dependency ^55,56^. UM represents such a malignancy, as recent transcriptomic and functional studies have demonstrated elevated autophagic activity and a strong reliance on lysosome-mediated degradation pathways for tumor maintenance and progression ^20,24,57^. Our findings further support this notion by demonstrating that pharmacological disruption of late-stage autophagy effectively suppresses UM cell survival *in vitro* (Fig. 5).

BetA has long been recognized for its anticancer activity and its ability to modulate autophagy in a highly context-dependent manner. Previous studies have reported that BetA can either induce autophagy initiation or inhibit autophagic flux, depending on cancer type, concentration, and cellular context ^29,32–36^. This duality of BetA-mediated autophagy regulation likely contributes to its antitumor efficacy but also limits its translational potential due to poor solubility, suboptimal pharmacokinetics, and limited target specificity ^37,58^. In the present study, we evaluated previously reported BetA-derivatives for their effects on UM cells and autophagy–lysosome function (Fig. 3). Among the BetA derivatives evaluated, ALS-997 emerged as a lead compound exhibiting potent autophagy-inhibitory activity, pronounced tumor selectivity, and robust antitumor effects (Fig. 5). Mechanistically, ALS-997 inhibited autophagy at a late stage by impairing lysosomal function rather than modulating upstream mTOR signaling (Fig. 6A, B). This distinction is particularly relevant, as inhibition of autophagosome degradation at the lysosomal level is increasingly recognized as a more effective strategy for disrupting autophagy-dependent cancer cell survival than targeting autophagy initiation alone. Our data identify NPC1 as a critical molecular target underlying ALS-997-mediated lysosomal dysfunction. NPC1 is an essential lysosomal transmembrane protein that regulates intracellular cholesterol trafficking and maintains lysosomal homeostasis (Fig. 7 and Fig. 8). Inhibition of NPC1 leads to lysosomal cholesterol accumulation, altered lysosomal positioning, impaired acidification, and blockade of autophagic flux (Fig. 6). These phenotypes closely mirror those observed following ALS-997 treatment, including lysosomal clustering, biphasic changes in Lysotracker staining (Fig. 6G, H), the accumulation of p62 and LC3-II (Fig. 6A, B). Notably, pharmacological and genetic inhibition of NPC1 has been reported to induce perinuclear accumulation and clustering of enlarged lysosomes, accompanied by increased Lysotracker fluorescence intensity, followed by loss of lysosomal acidity under conditions of severe dysfunction ^50,51^. The lysosomal phenotypes observed in ALS-997–treated cells are therefore highly consistent with previously reported NPC1 inhibition models. Importantly, Filipin staining demonstrated intracellular cholesterol accumulation upon ALS-997 treatment, comparable to that induced by the canonical NPC1 inhibitor U18666A ^59^, providing functional evidence for impaired cholesterol trafficking (Fig. 7A, B). Furthermore, CETSA analysis revealed that ALS-997 significantly increased the thermal stability of NPC1, supporting direct target engagement in a cellular context (Fig. 7C, D). Molecular docking analysis further suggested that ALS-997 interacts with NPC1 at the cholesterol-binding interface, forming stabilizing hydrogen bonds with key residues involved in sterol recognition (Fig. 8). Together, these complementary approaches strongly support a model in which ALS-997 directly inhibits NPC1, leading to lysosomal cholesterol sequestration and functional impairment. Disruption of NPC1-mediated lysosomal homeostasis had profound downstream consequences on cell fate. ALS-997 induced apoptotic signaling at substantially lower concentrations than BetA, as evidenced by caspase-3 activation, and PARP cleavage (Fig. 5). These findings are consistent with previous reports demonstrating that lysosomal dysfunction and autophagy blockade can sensitize cancer cells to apoptosis by exacerbating metabolic stress, oxidative damage, and proteotoxic burden ^60–62^. The pronounced tumor selectivity of ALS-997 suggests that UM cells are particularly vulnerable to NPC1-dependent lysosomal disruption, likely reflecting their heightened reliance on autophagy for survival. While higher dosing did not result in a strictly linear dose–response relationship, this observation may reflect complex pharmacodynamic effects of lysosomal perturbation in vivo and warrants further investigation.

In summary, this study demonstrates that targeting lysosomal cholesterol trafficking via NPC1 inhibition represents a promising strategy for disrupting autophagy-dependent survival in UM. A variety of autophagy inhibitors have been developed, including lysosomal inhibitors such as chloroquine and bafilomycin A1, as well as Vps34 inhibitors ^63,64^, ULK1/2 inhibitors ^65,66^, Atg4B inhibitors^67–70^, and compounds that disrupt formation of the Atg5-Atg12-Atg16L complex^71^; however, none of these agents have yet achieved successful clinical application. By overcoming the physicochemical and mechanistic limitations of BetA, ALS-997 serves as a BetA-derived NPC1-targeting compound that links lysosomal dysfunction, autophagy blockade, and antitumor activity. These findings support NPC1 as a potentially druggable regulator of autophagic flux in UM and provide a strong rationale for further development of NPC1-targeting therapeutics for UM and potentially other autophagy-dependent cancers.

## Conclusion

In this study, we demonstrated that BetA impairs autophagic flux in UM. By evaluating a series of previously reported BetA derivatives, we identified ALS-997 as a compound with substantially greater potency than BetA in UM cells and a strong ability to impair autophagic flux. ALS-997 exhibited selective cytotoxicity toward UM cells compared with normal melanocytes, and inhibited autophagic flux by inducing cholesterol accumulation and lysosomal dysfunction via NPC1 inhibition. Molecular docking analysis suggested that hydroxymethyl and phenolic moieties of ALS-997 may form hydrogen bonds with key NPC1 residues (Glu688 and Ile609), potentially contributing to its binding. ALS-997 inhibited autophagic flux and activated apoptotic signaling in UM 92.1 cells, resulting in pronounced anticancer activity. Collectively, these findings suggest that ALS-997 is a novel NPC1 inhibitor that effectively suppresses UM through autophagy inhibition, potentially representing a promising therapeutic strategy for enhancing treatment outcomes in UM.

## List of Abbreviations

AMPK: AMP-activated protein kinase
Atg: autophagy-related
BAP1: BRCA1-associated protein 1
BetA: betulinic acid
BSA: bovine serum albumin
CETSA: cellular thermal shift assay
CQ: chloroquine
FBS: Fetal Bovine Serum
GNAQ: G protein subunit alpha q
GNA11: G protein subunit alpha 11
IC_50_: half-maximal inhibitory concentration
LAMP2: lysosome-associated membrane protein 2
LC3: microtubule-associated protein 1 light chain 3
MAPK: mitogen-activated protein kinase
mTOR: mechanistic target of rapamycin
NPC1: Niemann-Pick disease type C1
PARP: poly ADP-ribose polymerase
PVDF: polyvinylidene fluoride
PBS: phosphate buffered saline
ROI: regions of interest
ROS: reactive oxygen species
SlogP: calculated partition coefficient
TAZ: transcriptional coactivator with PDZ-binding motif
tPSA: topological polar surface area
ULK1: Unc-51-like autophagy-activating kinase 1
UM: uveal melanoma
YAP: Yes-associated protein 1

## Declarations

## Ethics approval and concent to participate

This study did not involve human participants or animal experiments. Human-derived cells and DNA samples were used in accordance with institutional guidelines and ethical standards.

## Concent for publication

This study did not involve human participants.

## Availability of data and materials

The authors declare that the data supporting the findings of this studyare available within the paper and its Supporting Information. Furtherinformation and requests for resources and reagents should be directedto and will be fulfilled by the Lead Contact, Kaori H. Yamada.

## Authors’ contributions

Conceptualization, M.T.F. and K.H.Y., Project administration and supervision, K.H.Y., Data curation, Y.K., S.S., J.Y.N., G.L., and M.T.F., Formal analysis, Y.K., Methodology, Y.K., Investigation, Y.K., Visualization, Y.K., Validation, K.H.Y., Resources, M.T.F. and K.H.Y., Writing —original draft presentation, Y.K., Writing and editing, M.J.H., S.E., and K.H.Y. Funcing acquisition, Y.K., and K.H.Y., All authors have read and agreed to the submitted version of the manuscript.

## Acknowledgement

Portions of this work were carried out in the Fluorescent Imaging Core via the Research Resources Center (RRC) at the University of IllinoisChicago (UIC).

