## Supplemental Figures for "Development of Novel NPC1-Targeting Autophagy Inhibitors for the Treatment of Uveal Melanoma"

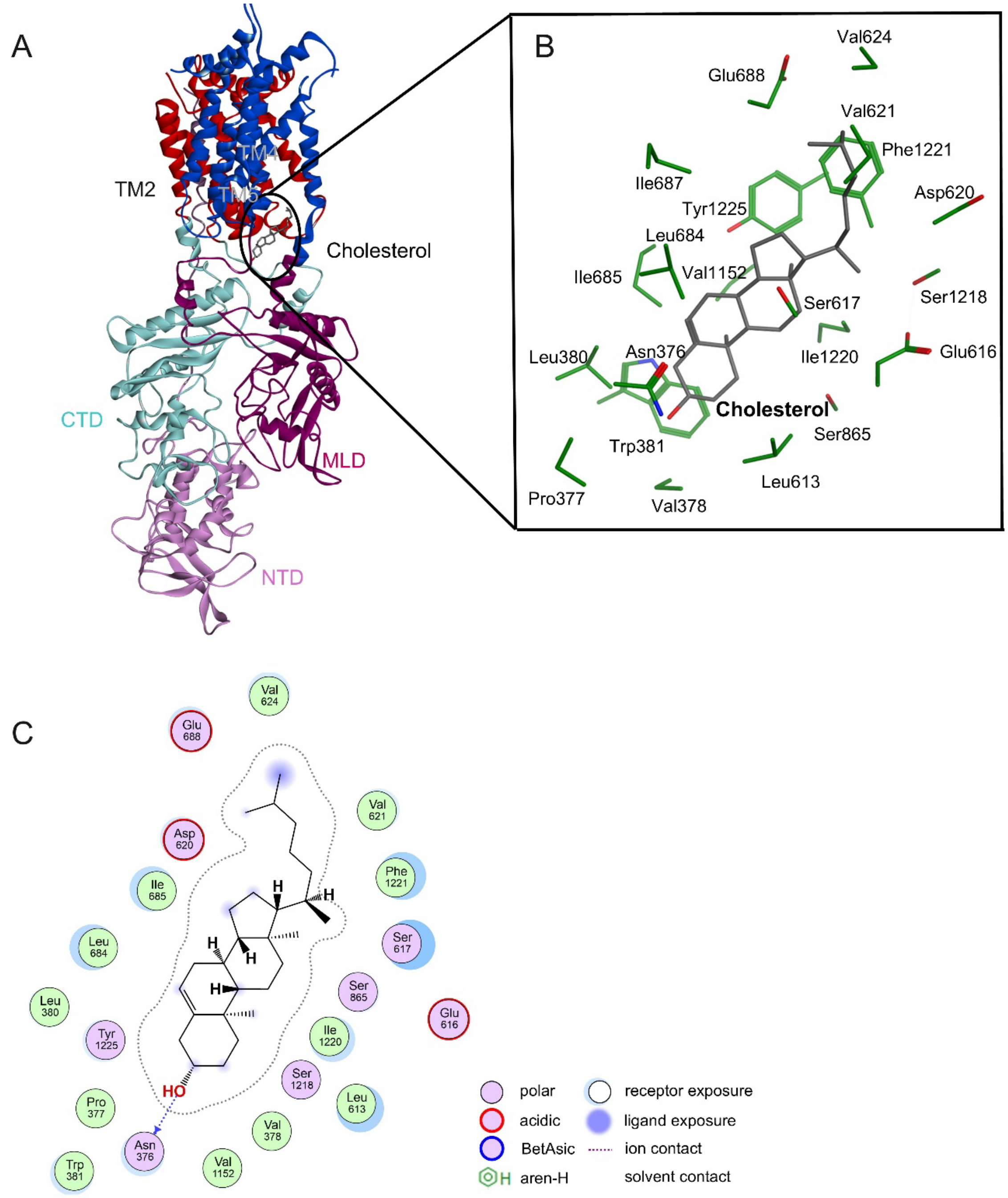


**Supplemental Figure S1. Docking model of NPC1 bound to cholesterol.**

(A) Overall docking model of cholesterol bound to human NPC1 structure [PDB: 6UOX]. NPC1 domains are shown in different colors: transmembrane helices/domain (TM), N-terminal luminal domain (NTD), middle luminal domain (MLD), and C-terminal luminal domain (CTD).

cholesterol is shown in gray. (B) Enlarged view of the predicted cholesterol binding pocket located near the interface between the MLD and the sterol-sensing domain-containing transmembrane region. Amino acid residues surrounding cholesterol are shown as sticks. (C) Two-dimensional ligand-protein interaction diagram of cholesterol with NPC1. The docking model predicts hydrogen-bond interactions between cholesterol and NPC1 residues, including Asn376. Docking calculations were performed using MOE2024.0601.

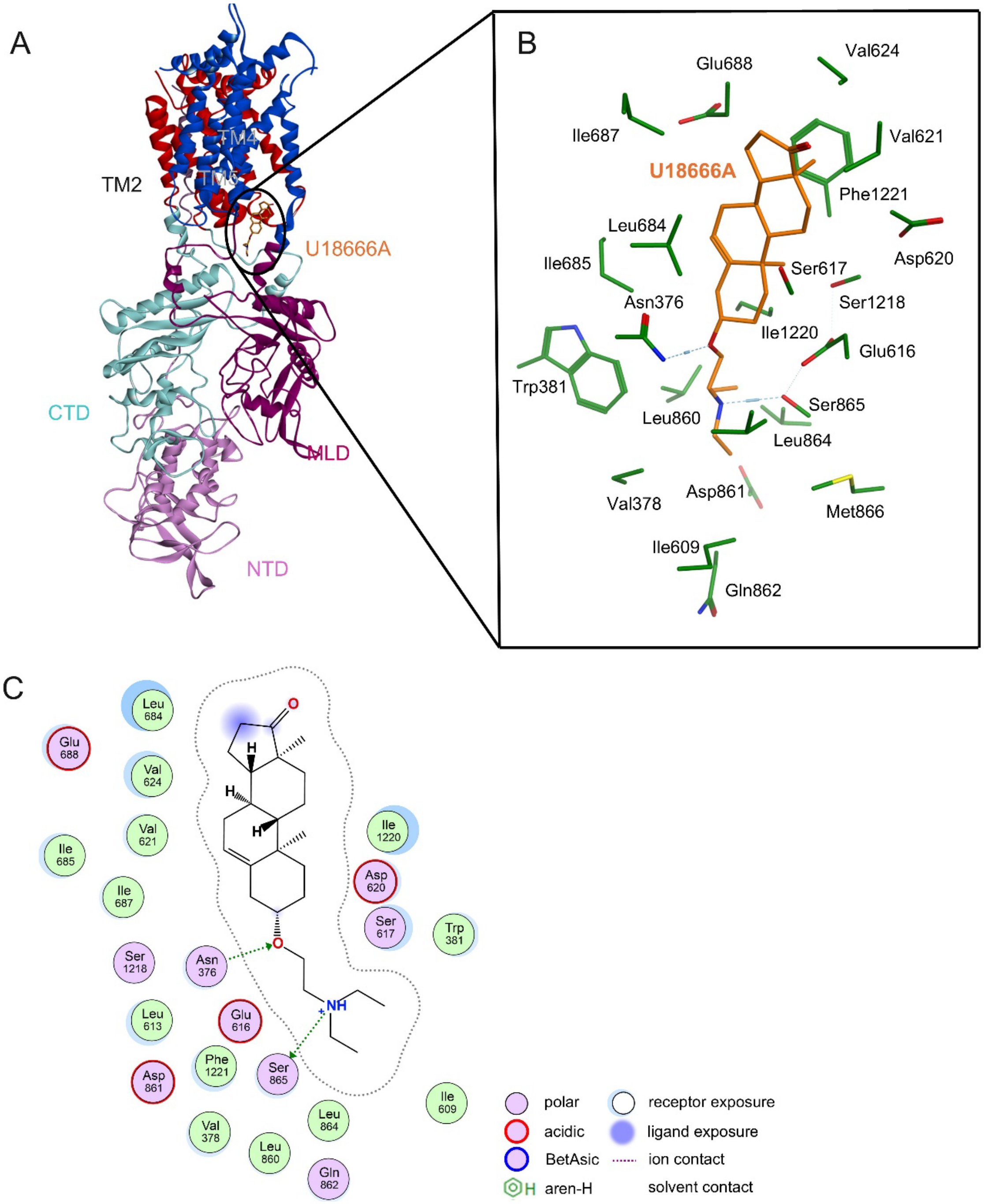


**Supplemental Figure S2. Docking model of NPC1 bound to U18666A.**

(A) Overall docking model of U18666A bound to human NPC1 structure [PDB: 6UOX]. U18666A is shown in orange. (B) Enlarged view of the predicted U18666A binding pocket located near the interface between the MLD and the sterol-sensing domain-containing transmembrane region.

Amino acid residues surrounding U18666A are shown as sticks. (C) Two-dimensional ligand-protein interaction diagram of U18666A with NPC1. The docking model predicts hydrogen-bond interactions between U18666A and NPC1 residues, including Asn376 and Ser865. Docking calculations were performed using MOE2024.0601.

**Table S1: The antibodies used for this study.**

| Target | Company | Catalog number |
|---|---|---|
| **Beclin 1** | Cell Signaling | 3738 |
| **Cleaved caspase 3** | Cell Signaling | 9664 |
| **GAPDH** | Santa Cruz | Sc-365062 |
| **LAMP2** | BD | 555803 |
| **LC3** | Proteintech | 14600-1-AP |
| **mTOR** | CUSABIO | CBP-PA008481 |
| **Phosphorylated mTOR** | CUSABIO | CBP-PA000652 |
| **NPC1** | Proteintech | 13926-1-AP |
| **p62** | COSMO BIO | MBL-PM045 |
| **PARP** | Cell Signaling | 9532 |
| **Cleaved PARP** | Cell Signaling | 5625 |
